# A Gatekept Deep Semiparametric Strategy for Evaluating Lung Transplant Treatment Effect Heterogeneity

**DOI:** 10.64898/2026.04.13.718254

**Authors:** Shuai Yuan, Fei Zou, Baiming Zou

**Affiliations:** Department of Biostatistics, University of North Carolina at Chapel Hill, Chapel Hill, North Carolina 27599, U.S.A.

**Keywords:** Cross-fitting, Deep learning, Heterogeneous treatment effects, Kernel score test, Lung transplantation, Permutation tests

## Abstract

Bilateral lung transplantation (BLT) can improve post-transplant lung function compared with single lung transplantation, but its additional functional benefit may not be uniform across recipients. Because BLT uses both donor lungs, transplant programs need to know whether its effect varies across clinically recognizable recipient profiles. Estimating such heterogeneity from observational registries is challenging. Donor and recipient characteristics influence both treatment selection and outcomes, and flexible individualized effect estimators can produce apparent subgroup variation even when no heterogeneity is present. We develop *deepHTL*, a gatekept semiparametric deep learning workflow that tests for heterogeneity, screens for the covariates driving it, and summarizes individualized and group average effects only when global evidence supports heterogeneity. *deepHTL* adjusts for confounding with cross-fitted bagged deep neural networks and separates the dominant average effect from residual heterogeneity through a revised estimator based on the Robinson transformation. In 14,196 adult recipients from the US national transplant registry, *deepHTL* finds strong evidence that the effect of BLT on forced expiratory volume in 1 second (FEV1) varies across recipients. Benefits are largest for recipients with low baseline FEV1, low lung allocation score, or low body mass index, whereas those with preserved baseline lung function gain little. Simulations under registry-like confounding show that *deepHTL* controls the type I error rate, identifies true effect modifiers, and estimates individualized effects accurately, supporting testing before interpretation in observational registries.

## 1 Introduction

For candidates awaiting lung transplantation, the choice between bilateral lung transplantation (BLT) and single lung transplantation (SLT) is clinically consequential and resource-sensitive. BLT can improve post-transplant lung function on average (De Oliveira et al., 2012; Force et al., 2011), but it uses both donor lungs and may impose greater surgical burden. Thus, the practical question for transplant programs is not only whether BLT is beneficial on average, but whether the additional functional benefit is large enough for particular recipient profiles to justify the use of two donor lungs (Munson et al., 2011). Post-transplant lung function is typically quantified by forced expiratory volume in 1 second (FEV1), a central measure of graft function that is closely related to long-term clinical prognosis (Verleden et al., 2019; Young et al., 2007). Heterogeneity in the FEV1 benefit of BLT is therefore of direct clinical relevance, and a useful analysis must answer three questions in order: whether BLT improves FEV1 on average, whether its effect varies across recipients, and, if it does, which donor or recipient characteristics drive the variation and how large the benefit is for clinically interpretable recipient profiles.

Lung transplant data usually come from observational registries, where treatment assignment is not random. If the resulting confounding is not rigorously addressed, it can substantially bias treatment effect estimates (Rigdon et al., 2018). Because many donor and recipient variables interact through nonlinear and nonadditive pathways, conventional treatment assignment and outcome models can be severely misspecified. Inferring causal heterogeneous treatment effects (HTE) from such data is therefore challenging. A range of machine learning methods has been developed to estimate covariate-dependent treatment effects, such as the T-, S- and X-learners (Künzel et al., 2019), causal forests (Wager and Athey, 2018) and Bayesian causal forests (Hahn et al., 2020). These methods can, however, be sensitive to limited overlap and to the inductive bias of the base learner (D’Amour et al., 2021; Curth and van der Schaar, 2021; Lu et al., 2017). Semiparametric regression based on the Robinson transformation (Robinson, 1988) offers a robust alternative. It applies Neyman-orthogonal estimating equations to mitigate the bias from estimating the nuisance functions, and Nie and Wager (2021) formalized this approach for HTE estimation as the quasi-oracle R-learner. Nevertheless, the robustness of the R-learner relies heavily on the quality of the nuisance estimators. When the treatment effect is large, the residuals become bimodal even when no HTE is present, which biases the nuisance estimates (Mi et al., 2019), and these errors propagate into the HTE estimates (Chernozhukov et al., 2018).

The methods above focus only on estimating individualized effects. However, in a flexible machine learning analysis, a spread of estimated treatment effects can appear even when the true treatment effect is constant. Without a formal global test, this apparent variation cannot be distinguished from sampling variability or nuisance estimation error, and residual confounding further complicates its causal interpretation. Testing for HTE should therefore precede estimation and subgroup interpretation, yet formal tests remain limited. Existing procedures use series approximations (Crump et al., 2008), focus on randomized experiments (Chang et al., 2021) or examine prespecified subgroups (Dai and Stern, 2022). Beyond a global test, a transplant program also needs to know which recipient characteristics drive the heterogeneity. Prespecified subgroup comparisons examine only the modifiers that were anticipated, and variable importance rankings do not by themselves provide calibrated covariate-specific tests. None of these approaches offers the integrated sequence of testing, screening and estimation that the registry analysis requires.

To meet this need, we develop *deep Heterogeneous Treatment Learning* (*deepHTL*) as a gatekept analysis strategy for observational HTE studies. The workflow first tests whether treatment effect heterogeneity exists, then screens for the covariates that drive it, and summarizes individualized and group average treatment effects only after the global test supports heterogeneity. The strategy is designed for comparative effectiveness studies of registry data, in which treatment assignment is confounded and clinical interpretation requires both valid inference and interpretable effect modifiers. A bagged deep neural network (bagged-DNN) estimates the nuisance functions under complex confounding, and a revised estimator based on the Robinson transformation reduces their bias when the average effect is large.

We apply *deepHTL* to 14,196 adult lung transplant recipients in the United Network for Organ Sharing (UNOS) registry to evaluate the effect of BLT versus SLT on post-transplant FEV1. The analysis finds strong evidence of treatment effect heterogeneity, with baseline FEV1, lung allocation score and recipient body mass index (BMI) among the covariates that modify the effect. The estimated BLT benefit is largest among recipients with low baseline FEV1, low lung allocation score or low BMI, whereas recipients with preserved baseline lung function show limited FEV1 benefit. Targeted simulations then show why the revised estimator is needed. When the average treatment effect is large, removing the constant component improves the fit of the outcome model and keeps both global tests near their nominal level.

## 2 Motivating UNOS lung transplant study

We investigate treatment effect heterogeneity using the UNOS lung transplant registry. The analytic cohort consists of 14,196 adult recipients transplanted between May 2005 and September 2011. The treatment indicator is *Z* = 1 for BLT and *Z* = 0 for SLT, with 9,678 BLT recipients and 4,518 SLT recipients. The outcome *Y* is the recipient’s FEV1 measured one year after transplantation (on average 379 days), expressed as a percentage of the value predicted for a healthy person of the recipient’s age, sex, height and race (Stanojevic et al., 2022) and denoted FEV1%.

The observational nature of the registry creates the central statistical challenge. BLT and SLT recipients differed in observed characteristics (Table S1). BLT recipients were younger (53.4 versus 63.4 years), had lower BMI (24.8 versus 26.3) and lower baseline FEV1% (37.1 versus 43.6), and more often required ventilator support (4.8% versus 1.6%). Because these characteristics are related both to the choice of procedure and to post-transplant lung function, an unadjusted comparison of BLT and SLT recipients is subject to confounding by indication. The 26 covariates in ***X*** are those selected by the group lasso doubly robust (GL-DR) procedure (Koch et al., 2018). These covariates are mostly weakly correlated, with a median absolute pairwise correlation of 0.03 and only 23 of the 325 pairs above 0.2 in absolute value (Figure S1). Four pairs exceed 0.5 in absolute value, the strongest being recipient height with recipient sex (|*r*| = 0.70), which matters when heterogeneity is attributed to individual covariates (Section 5). The causal estimand is the conditional average treatment effect, the expected difference between the potential post-transplant FEV1% under BLT and under SLT given ***X***. When the selection and outcome mechanisms are too complex for conventional regression models, inadequate adjustment for confounding leaves a bias in the estimated effect that varies with ***X***, which can produce spurious heterogeneity. This motivates the sequence of testing, screening and estimation that the next section develops as *deepHTL*.

## 3 Methods

### 3.1 Analysis workflow and assumptions

*deepHTL* is a gatekept analysis in five steps. First, the propensity score and the outcome mean are estimated by cross-fitted bagged-DNNs. Second, the constant component of the treatment effect is removed by the revised transformation. Third, the global null hypothesis of a constant effect is tested. Fourth, if it is rejected, the candidate covariates are screened for effect modification. Fifth, individualized and group average effects are estimated and interpreted in light of the screen.

Let *Y*_*i*_(1) and *Y*_*i*_(0) denote the potential FEV1% outcomes of recipient *i* under BLT (*Z*_*i*_ = 1) and SLT (*Z*_*i*_ = 0), and *τ* (***X***) = E{*Y* (1) − *Y* (0) | ***X***} the conditional average treatment effect. The assumptions include:

i. consistency, *Y*_*i*_ = *Z*_*i*_*Y*_*i*_(1) + (1 − *Z*_*i*_)*Y*_*i*_(0),
ii. conditional exchangeability (no unmeasured confounding), {*Y*_*i*_(1), *Y*_*i*_(0)} **╨** *Z*_*i*_ | ***X***_*i*_,
iii. overlap, *η* ≤ Pr(*Z*_*i*_ = 1 | ***X***_*i*_) ≤ 1 − *η* almost surely for some *η* ∈ (0, 1*/*2).

### 3.2 Deep heterogeneous treatment learning for estimating HTE

To represent potentially complex HTE while leaving the confounding structure unspecified, we consider the semiparametric regression model

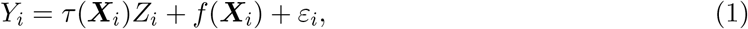

where the nuisance function *f* (***X***) and the conditional average treatment effect *τ* (***X***) are left unspecified and E (*ε*_*i*_ | ***X***_*i*_, *Z*_*i*_) = 0 with finite variance. To avoid estimating *f* (***X***) in (1), we apply the Robinson transformation

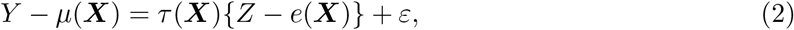

where the nuisance functions are the conditional mean outcome *µ*(***X***) = E[*Y* | ***X***] and the propensity score *e*(***X***) = E[*Z* | ***X***]. They can be estimated by kernel methods (Robinson, 1988) or by machine learning methods such as the least absolute shrinkage and selection operator (Lasso) (Tibshirani, 1996), extreme gradient boosting (XGBoost) (Chen and Guestrin, 2016) and kernel ridge regression (KRR) (Vovk, 2013), as in Nie and Wager (2021). These learners may, however, be inadequate for nonlinear and nonadditive confounding structures, in which case the nuisance estimates can be systematically biased in finite samples, and this bias propagates into the estimate of *τ* (***X***) (Chernozhukov et al., 2018). A deep neural network (DNN) can approximate a wide class of functions to an arbitrary level of accuracy (Hornik et al., 1989; Cybenko, 1989), but its finite-sample performance can be poor because training from random initial parameters may not reach the global optimum. The bagged-DNN method addresses this problem by aggregating predictions from multiple DNNs trained on bootstrap samples and discarding poorly performing networks, which stabilizes the fit against the randomness of initialization. We therefore adopt the bagged-DNN for estimating the nuisance functions (Mi et al., 2019, 2021; Zou et al., 2025).

To reduce overfitting bias in the nuisance estimates, we randomly partition the observation indices {1, … , *n*} into *K* disjoint folds *I*_1_, … , *I*_*K*_, with *K* = 5 in all analyses. For each fold *k*, the bagged-DNN models for *µ*(***X***) and *e*(***X***) are trained on the observations outside *I*_*k*_ and predict only for the held-out observations *i* ∈ *I*_*k*_. We then estimate *τ* (***X***) by minimizing the squared loss derived from (2) with these out-of-fold nuisance estimates 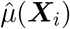 and 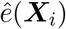:

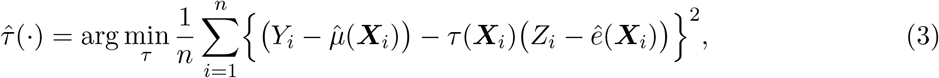

where the minimization is carried out with the bagged-DNN. However, when treatment effects are large relative to outcome noise, the finite-sample accuracy of this estimator can deteriorate. By (2), the residual *Y* − *µ*(***X***) is the sum of *τ* (***X***){*Z* − *e*(***X***)} and the noise *ε*. When |*τ* (***X***)| is small, the residual is dominated by *ε*, whose distribution is typically unimodal and well suited to least squares estimation. When |*τ* (***X***)| is large, *τ* (***X***){*Z* −*e*(***X***)} dominates and the residual distribution becomes bimodal, which biases 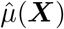 in finite samples, and this bias propagates to 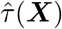 (Mi et al., 2019). We therefore decompose the treatment effect into a constant average component and a residual heterogeneous component, *τ* (***X***) = *τ*_0_ + *r*(***X***), and reformulate (2) as

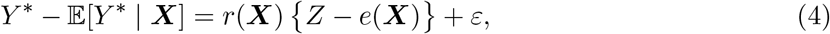

where *Y* ^*^ = *Y* −*τ*_0_*Z* is the centered outcome and *µ*^*^(***X***) = E(*Y* ^*^ | ***X***) is estimated with the bagged- DNN under the same cross-fitting scheme. Centering removes the constant component from the residual, which is then close to unimodal, so that *µ*^*^(***X***) can be estimated more accurately. We estimate *τ*_0_ by the computationally efficient Neyman-orthogonal estimator:

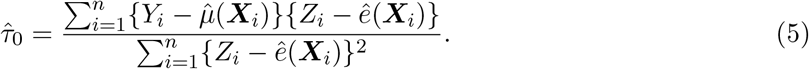

The estimator 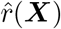 of the residual heterogeneous component is then obtained by minimizing the squared loss

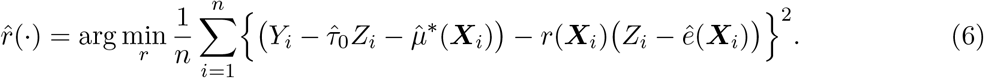

The revised estimator of *τ* (***X***) combines the two components,

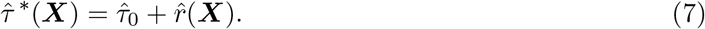

### 3.3 Testing for HTE

Before any individualized effect or subgroup is interpreted, we test the null hypothesis *H*_0_ : Var{*τ* (***X***)} = 0, under which *τ* (***X***) equals an unknown constant. We propose two global tests, a computationally efficient kernel score test and a more robust but computationally intensive permutation test, and extend the permutation framework to covariate-specific tests that identify the sources of heterogeneity once the global null is rejected.

#### Kernel score test

The score test is based on residuals relative to the constant effect, so it requires an accurate estimate of this constant to distinguish heterogeneity from estimation error. The estimator 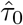 in (5), hereafter the unrevised estimator, provides an initial estimate, but its accuracy can deteriorate in finite samples when the treatment effect is large (Section 3.2). We therefore compute the revised estimator

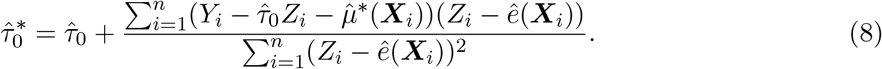

Using 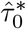, we build the score test from the partialling out score of the partially linear model (Chernozhukov et al., 2018). Let 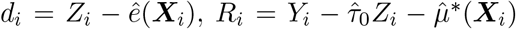 and 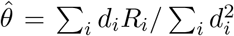. The score 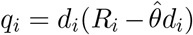 measures the outcome’s deviation from the fitted constant effect model, weighted by the treatment residual *d*_*i*_. Following the variance component approach (Liu et al., 2007; Lin et al., 2013), we use the kernel score statistic

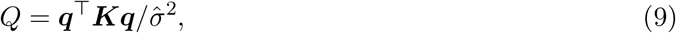

where 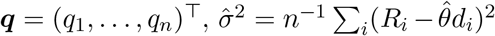 and ***K*** is a symmetric positive semidefinite kernel matrix with entries *K*_*ij*_ = *K*(***X***_*i*_, ***X***_*j*_) for a Mercer kernel *K*(·, ·), here the Gaussian radial basis function with bandwidth set to the median pairwise distance between standardized covariate vectors. Under *H*_0_ and regularity conditions, we approximate the null distribution of *Q* by 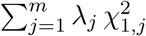, where 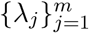 are the nonzero eigenvalues of ***P DKDP*** , with ***d*** = (*d*_1_, … , *d*_*n*_)^⊤^, ***D*** = diag(***d***) and ***P*** = ***I***_*n*_ − ***dd***^⊤^*/*(***d***^⊤^***d***). The analytic *p*-value 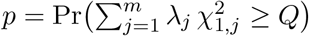 is computed by the Davies method (Davies, 1980; Wu et al., 2011). Technical details are given in Appendix C.

#### Permutation test

The kernel score test is powerful and computationally efficient, but its null distribution rests on a large-sample approximation that can be inaccurate in finite samples (Chen et al., 2016). We therefore also propose a permutation test that constructs the null distribution empirically.

We construct the test statistic from the out-of-sample prediction loss. Using the same *K* folds as in the nuisance estimation stage, we cross-fit the residual component model (6), so that the predictions 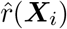 for *i* ∈ *I*_*k*_ come from a model trained on the remaining folds, and compute

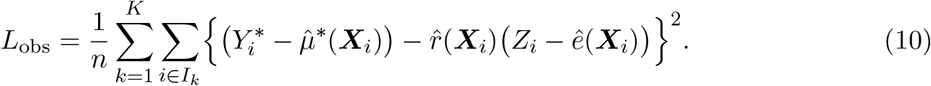

For *b* = 1, … , *B* we randomly permute the indices of the held-out predictions within each fold and treatment arm. With *π*^(*b*)^(*i*) denoting the permuted index of observation *i*,

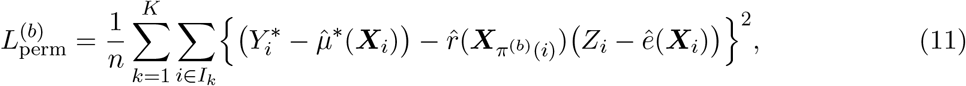

and 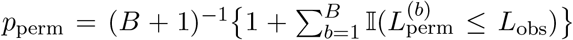. Under *H*_0_, the treatment effect is constant across covariate values. When the fitted model captures heterogeneity, permuting its predictions disrupts their alignment with the observations and is expected to increase the loss. A small *p*_perm_ indicates that the original predictions fit the observations better than the permuted predictions. For the unrevised estimator 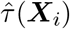, we use the same procedure with the cross-fitted loss in (3). Appendix A gives the complete estimation and testing algorithms. The covariate-specific screen applies this loss comparison to one covariate at a time. For each candidate *x*_*j*_, we permute its values within each held-out fold while keeping the fitted models, nuisance estimates and other covariates fixed. We then reevaluate 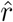 at the permuted covariates and compare the resulting losses with *L*_obs_.

## 4 Analysis of the UNOS lung transplant registry

We now apply *deepHTL* to the UNOS cohort of Section 2 to assess whether the BLT effect varies across recipients and clinically relevant subgroups.

### 4.1 Testing whether the BLT effect is heterogeneous

The global test is the gatekeeping step. Both the permutation test and the analytic kernel score test reject the null hypothesis of a constant BLT effect (*p <* 0.001 for both).

### 4.2 Identifying covariates that drive heterogeneity

Applied to all 26 adjustment covariates, the covariate-specific screen rejects the null hypothesis of no effect modification for eight of them at a false discovery rate of 5% (Benjamini and Hochberg, 1995) (Table 1). FEV1 is a prognostic marker of mortality (Young et al., 2007), the lung allocation score reflects clinical urgency and expected transplant benefit (Egan et al., 2006), and recipient obesity is associated with a failure to reach normal lung function after BLT (Keller et al., 2024). Correlation among the covariates requires attention in interpreting the remaining selections. Recipient height is correlated with recipient sex (|*r*| = 0.70, Section 2) and with donor height (0.55), the oxygen requirement at rest with the lung allocation score (0.49), and recipient age with recipient BMI (0.34) and baseline FEV1% (0.31). For such correlated covariates, the screen may not fully distinguish individual effect modifiers (Section 5.1, Table 4). These covariates are nevertheless clinically relevant. Height and sex inform lung size matching, and a higher donor-to-recipient predicted lung capacity ratio has been associated with better survival after BLT (Eberlein et al., 2013). The oxygen requirement at rest is an input to the lung allocation score (Egan et al., 2006), and the reported survival advantage of BLT is smaller in older recipients (Shacker et al., 2025). Recipient Hispanic ethnicity is essentially uncorrelated with the other covariates (|*r*| ≤ 0.14), but its selection rests on 982 recipients and is one of the two weakest (unadjusted *p* = 0.010). It may be a false discovery not ruled out at the 5% false discovery rate, or may reflect primary diagnosis, which is not among the covariates and differs between Hispanic and non-Hispanic White recipients (Bonner et al., 2023). We use the screen to guide subgroup interpretation rather than choosing modifiers from clinically appealing subgroup estimates.

**Table 1:** Covariate-specific screen on the UNOS cohort (*n* = 14,196) over the 26 adjustment covariates, ordered by *p. p*_BH_ is the Benjamini–Hochberg adjusted *p*-value. CMV denotes cytomegalovirus, TBI traumatic brain injury and PO_2_ the partial pressure of oxygen.

| Rank | Covariate | $p$ | $p_{BH}$ |
| --- | --- | --- | --- |
| 1 | Baseline FEV1% | <0.001 | 0.003 |
| 2 | Lung allocation score | <0.001 | 0.003 |
| 3 | Recipient BMI | <0.001 | 0.003 |
| 4 | Recipient height | <0.001 | 0.003 |
| 5 | Donor height | <0.001 | 0.003 |
| 6 | Oxygen requirement at rest | 0.001 | 0.006 |
| 7 | Recipient Hispanic ethnicity | 0.010 | 0.036 |
| 8 | Recipient age | 0.011 | 0.036 |
| 9 | Donor sex | 0.022 | 0.065 |
| 10 | Ischemic time | 0.028 | 0.073 |
| 11 | Donor cause of death, TBI | 0.113 | 0.268 |
| 12 | Donor to recipient distance | 0.184 | 0.371 |
| 13 | Six-minute walk distance | 0.185 | 0.371 |
| 14 | Local or regional allocation | 0.252 | 0.469 |
| 15 | Recipient sex | 0.297 | 0.494 |
| 16 | Donor diabetes | 0.304 | 0.494 |
| 17 | Race match | 0.365 | 0.559 |
| 18 | Ventilator support | 0.451 | 0.652 |
| 19 | Time to follow-up FEV1 | 0.522 | 0.714 |
| 20 | Donor CMV serology | 0.613 | 0.791 |
| 21 | Recipient diabetes | 0.639 | 0.791 |
| 22 | Donor BMI | 0.759 | 0.870 |
| 23 | Recipient $PO_2$ | 0.770 | 0.870 |
| 24 | Donor age | 0.810 | 0.878 |
| 25 | Sex match | 0.884 | 0.919 |
| 26 | Donor race, Black | 0.970 | 0.970 |

### 4.3 Estimating individualized and group average BLT benefit

Because the global test rejects homogeneity, we estimate *τ* (***X***) with the *deepHTL* estimator (7) and refer to 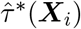 as the estimated individualized treatment effect (ITE) of recipient *i*. The estimated average treatment effect of BLT is 17.98 percentage points on the FEV1% scale (95% confidence interval 17.23 to 18.73), but the distribution of estimated ITEs (Figure 1) shows that this average masks substantial heterogeneity. The estimates span a wide range (−1.5 to 40.4 percentage points) and are distinctly bimodal. The main clinical message is therefore not simply that BLT improves FEV1% on average, but that the magnitude of improvement differs sharply across recipient profiles.

**Figure 1:**
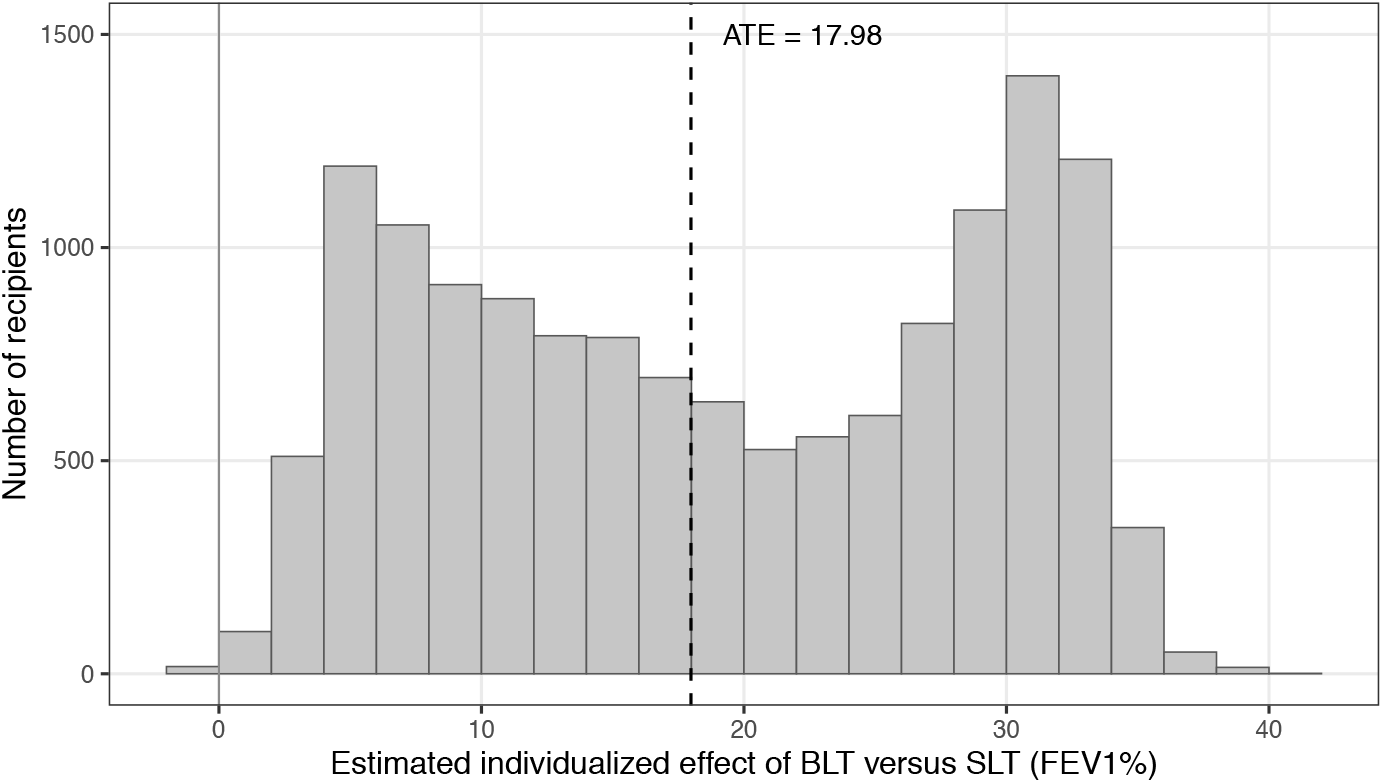
Estimated individualized treatment effects (BLT versus SLT) on post-transplant FEV1% in the UNOS cohort. The dashed line marks the estimated average treatment effect (17.98 percentage points).

To summarize the individualized estimates for clinically interpretable subgroups, we estimate group average treatment effects (GATE) using prespecified thresholds or categories for the eight selected covariates (Figure 2, Table S2). The estimated advantage of BLT is greatest in the sub-groups with baseline FEV1% below 30, lung allocation score below 35 and BMI below 18.5, and is substantially attenuated among recipients whose baseline FEV1% exceeds 50. It also declines with recipient height, from 21.1 percentage points below 165 cm to 15.5 at 175 cm or above. The panels of the other selected covariates call for more caution. The advantage is larger with donors of 175 cm or taller (20.1 versus 16.1 and 16.5 with shorter donors), although it falls with recipient height, with which donor height is positively correlated. Recipient sex was not selected, although the advantage is larger in women than in men (20.9 versus 16.1, Table S2). Because sex is strongly correlated with recipient height, the fitted model may use height as a proxy for sex, and the screen may not fully distinguish their individual contributions. The advantage is smaller when the oxygen requirement at rest exceeds 4 L/min (14.1 versus 20.1 and 20.9 at lower requirements), and this persists within the strata of baseline FEV1% below 30 and from 30 to 50. It is also smaller for Hispanic recipients (14.5 versus 18.2), but only among those with baseline FEV1% below 30 (21.0 versus 30.9) and not in the other two strata. Finally, it declines with recipient age, from 21.7 below 50 years to 12.0 at 70 to 74 and 11.7 at 75 or older, but the decline disappears within the strata of baseline FEV1%, suggesting that the age pattern may partly reflect differences in baseline lung function. Overall, GATE at prespecified thresholds or categories generally agree with the screen, and the other prespecified subgroups vary little apart from recipient sex (Table S2). The two analyses are complementary, as the screen provides covariate-level evidence of heterogeneity and GATE show the direction and size of subgroup differences.

**Figure 2:**
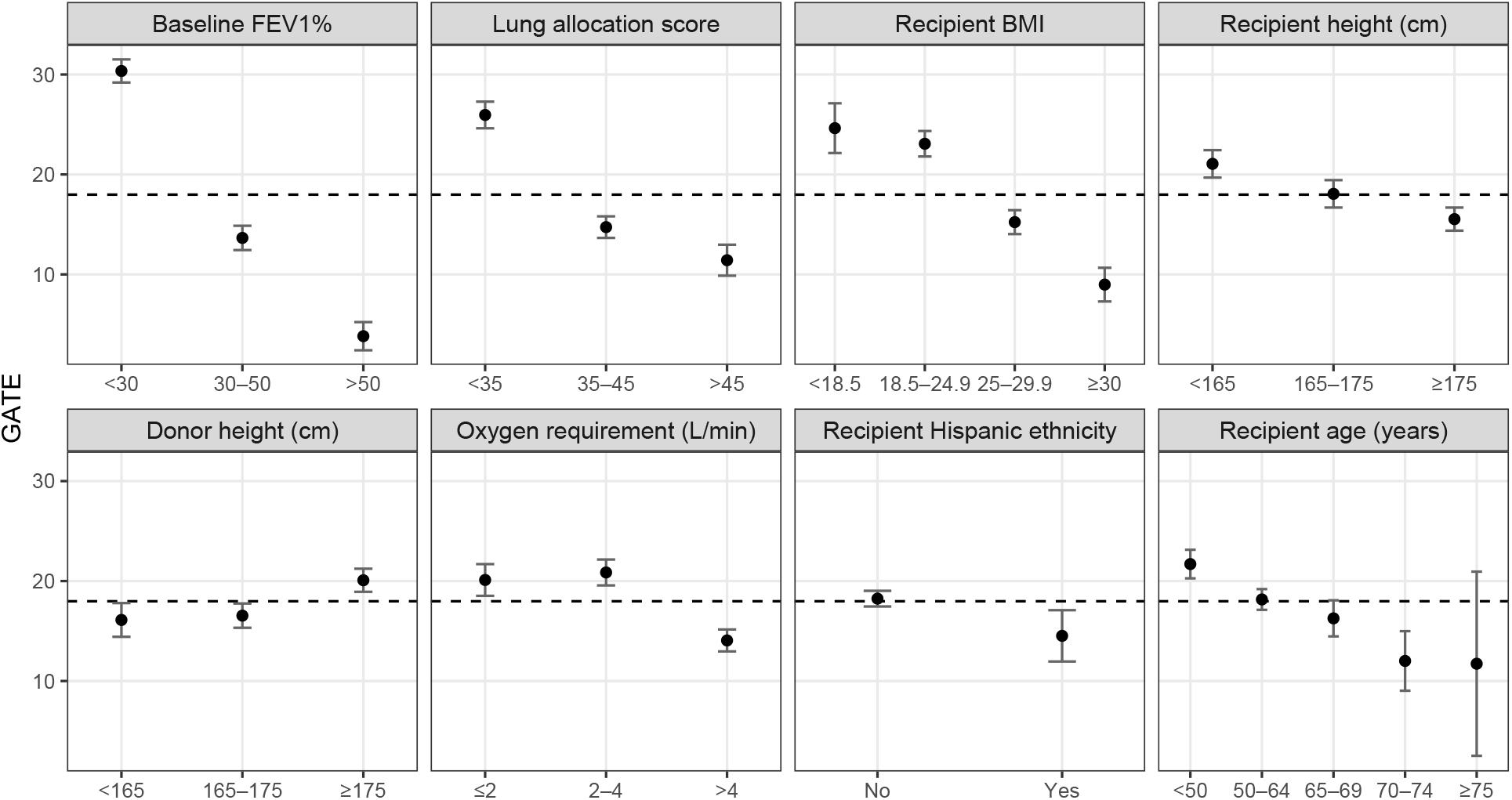
Group average treatment effects (BLT versus SLT) on post-transplant FEV1%, with 95% confidence intervals, for the eight covariates selected by the covariate-specific screen. The horizontal dashed line denotes the average treatment effect.

In summary, the additional functional benefit of BLT is most clearly concentrated among recipients with poorer baseline lung function, lower lung allocation score and lower BMI, and may also vary with donor and recipient height and with the oxygen requirement at rest. These findings do not by themselves constitute an allocation rule, but they show how a gatekept HTE analysis can identify recipient profiles for which the additional functional benefit of BLT appears most pronounced.

## 5 Targeted simulation studies

The simulations address four questions raised by the UNOS analysis: (i) whether the global tests control the type I error under a constant treatment effect, zero or large, with and without the revised estimator, (ii) how much power they have against nonlinear heterogeneity, (iii) whether the covariate-specific screen detects the true modifiers while keeping the type I error rate for the null covariates near the nominal level, and (iv) how accurately the revised estimator recovers *τ* (***X***) relative to established methods. Data are generated from the outcome model

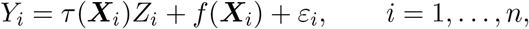

where ***X***_*i*_ denotes the *p*-dimensional covariate vector, including both confounders and null covariates, and *ε*_*i*_ ∼ *N* (0, *σ*^2^) is independent noise. Binary treatment assignment follows

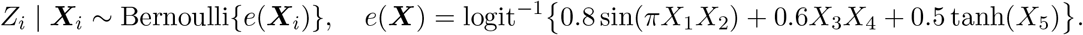

The nuisance function is 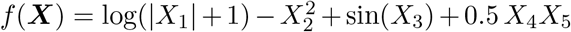, and the treatment effect function *τ* (***X***) varies by experiment:

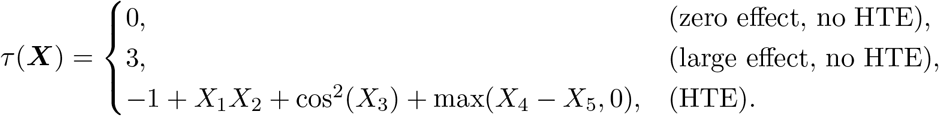

The covariates mimic the mix of continuous and binary variables and the correlation found in observational lung transplant data. We draw ***W***_*i*_ ∼ *N*_*p*_(**0, Σ**) with cor(*W*_*j*_, *W*_*k*_) = *ρ* for 1 ≤ *j*≠ *k* ≤ 10 and 0 otherwise, dichotomize three components, *X*_3_ = *N*{*W*_3_ *>* Φ^−1^(0.6)}, *X*_6_ = I{*W*_6_ *>* Φ^−1^(0.6)} and *X*_7_ = I{*W*_7_ *>* Φ^−1^(0.5)}, and set *X*_*j*_ = *W*_*j*_ for the remaining continuous covariates. We evaluate performance over a grid of block correlations *ρ* ∈ {0, 0.3}, sample sizes *n* ∈ {1000, 2000}, covariate dimensions *p* ∈ {20, 40} and noise levels *σ* ∈ {1, 3}, using 1000 independent replicates per configuration.

### 5.1 Performance on testing HTE

We first evaluate the two global tests under the null hypothesis of a homogeneous treatment effect, with *τ* = 0 and with *τ* = 3. Both tests are computed with the unrevised and with the revised estimator, and we report empirical type I error rates at the nominal level *α* = 0.05 (Table 2). Under *τ* = 0 the rejection rates of both the unrevised and the revised tests are close to the nominal level, except for the analytic test at *n* = 1000, *p* = 40, *σ* = 1 and *ρ* = 0.3, where the nuisance estimates are less accurate. This inflation disappears as the sample size increases. Under *τ* = 3 the unrevised analytic test fails to control the type I error at low noise, with rejection rates up to 0.098, and the unrevised permutation test reaches up to 0.097 in the same configurations. This is the failure mode anticipated in Section 3.2. When |*τ*_0_| is large relative to *σ*, the resulting bimodality in the residual *Y* − *µ*(***X***) biases 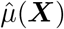, and that bias propagates to the tests as spurious heterogeneity. The revised estimator restores type I error control by removing the constant component before the nuisance functions are estimated, and both revised tests are close to the nominal level in every configuration.

**Table 2:** Empirical type I error rates at the nominal level *α* = 0.05 for the analytic kernel score test (Anal.) and the permutation test (Perm.), using the unrevised and the revised estimator, under the registry design at block correlation *ρ* = 0 and 0.3. Bold entries indicate inflated type I error.

| $\tau$ | Estimator | $n$ | $p$ | $\sigma$ | $\rho = 0$ | | $\rho = 0.3$ | |
| --- | --- | --- | --- | --- | --- | --- | --- | --- |
|  |  |  |  |  | Anal. | Perm. | Anal. | Perm. |
| $\tau = 0$ | Unrevised | 1000 | 20 | 1 | 0.047 | 0.053 | 0.053 | 0.055 |
|  |  | 1000 | 20 | 3 | 0.053 | 0.057 | 0.056 | 0.055 |
|  |  | 1000 | 40 | 1 | 0.045 | 0.050 | **0.081** | 0.065 |
|  |  | 1000 | 40 | 3 | 0.047 | 0.050 | 0.054 | 0.063 |
|  |  | 2000 | 20 | 1 | 0.050 | 0.048 | 0.062 | 0.067 |
|  |  | 2000 | 20 | 3 | 0.047 | 0.054 | 0.060 | 0.056 |
|  |  | 2000 | 40 | 1 | 0.048 | 0.057 | 0.051 | 0.059 |
|  |  | 2000 | 40 | 3 | 0.054 | 0.064 | 0.060 | 0.052 |
|  | Revised | 1000 | 20 | 1 | 0.056 | 0.058 | 0.047 | 0.056 |
|  |  | 1000 | 20 | 3 | 0.053 | 0.057 | 0.056 | 0.048 |
|  |  | 1000 | 40 | 1 | 0.048 | 0.044 | 0.071 | 0.065 |
|  |  | 1000 | 40 | 3 | 0.045 | 0.051 | 0.049 | 0.063 |
|  |  | 2000 | 20 | 1 | 0.051 | 0.054 | 0.068 | 0.063 |
|  |  | 2000 | 20 | 3 | 0.051 | 0.050 | 0.058 | 0.060 |
|  |  | 2000 | 40 | 1 | 0.047 | 0.050 | 0.049 | 0.068 |
|  |  | 2000 | 40 | 3 | 0.052 | 0.068 | 0.053 | 0.058 |
| $\tau = 3$ | Unrevised | 1000 | 20 | 1 | **0.087** | **0.097** | **0.086** | **0.092** |
|  |  | 1000 | 20 | 3 | 0.055 | 0.057 | 0.064 | 0.065 |
|  |  | 1000 | 40 | 1 | **0.079** | **0.076** | **0.092** | **0.082** |
|  |  | 1000 | 40 | 3 | 0.049 | 0.060 | 0.060 | 0.059 |
|  |  | 2000 | 20 | 1 | **0.095** | **0.081** | **0.080** | **0.093** |
|  |  | 2000 | 20 | 3 | 0.053 | 0.048 | 0.049 | 0.053 |
|  |  | 2000 | 40 | 1 | **0.098** | **0.090** | **0.080** | **0.084** |
|  |  | 2000 | 40 | 3 | 0.048 | 0.066 | 0.053 | 0.062 |
|  | Revised | 1000 | 20 | 1 | 0.053 | 0.054 | 0.041 | 0.053 |
|  |  | 1000 | 20 | 3 | 0.050 | 0.059 | 0.060 | 0.062 |
|  |  | 1000 | 40 | 1 | 0.038 | 0.052 | 0.053 | 0.052 |
|  |  | 1000 | 40 | 3 | 0.052 | 0.056 | 0.055 | 0.060 |
|  |  | 2000 | 20 | 1 | 0.049 | 0.051 | 0.050 | 0.051 |
|  |  | 2000 | 20 | 3 | 0.050 | 0.050 | 0.047 | 0.056 |
|  |  | 2000 | 40 | 1 | 0.050 | 0.051 | 0.062 | 0.068 |
|  |  | 2000 | 40 | 3 | 0.046 | 0.061 | 0.057 | 0.064 |

Table 3 shows that the revised tests do not control the type I error at the expense of power. Under the alternative, both tests have power above 0.9 in every configuration with *σ* = 1, and at *σ* = 3 power increases markedly with *n*. In the more challenging configurations with *σ* = 3, the permutation test is more powerful than the analytic test in all but one of them, and correlation among the covariates costs the analytic test far more power than the permutation test.

**Table 3:** Empirical power of the global tests for detecting HTE using the revised estimator, at the nominal level *α* = 0.05, for the analytic kernel score test (Anal.) and the permutation test (Perm.) under the registry design at block correlation *ρ* = 0 and 0.3.

| $n$ | $p$ | $\sigma$ | $\rho = 0$ | | $\rho = 0.3$ | |
| --- | --- | --- | --- | --- | --- | --- |
|  |  |  | Anal. | Perm. | Anal. | Perm. |
| 1000 | 20 | 1 | 1.000 | 1.000 | 1.000 | 1.000 |
| 1000 | 20 | 3 | 0.647 | 0.672 | 0.430 | 0.623 |
| 1000 | 40 | 1 | 1.000 | 1.000 | 0.991 | 0.991 |
| 1000 | 40 | 3 | 0.469 | 0.427 | 0.256 | 0.296 |
| 2000 | 20 | 1 | 1.000 | 1.000 | 1.000 | 1.000 |
| 2000 | 20 | 3 | 0.958 | 0.985 | 0.843 | 0.977 |
| 2000 | 40 | 1 | 1.000 | 1.000 | 1.000 | 1.000 |
| 2000 | 40 | 3 | 0.880 | 0.945 | 0.601 | 0.908 |

For the covariate-specific screen, *X*_1_ to *X*_5_ are effect modifiers and *X*_6_ to *X*_20_ are null covariates (Table 4). At both correlation levels the screen recovers the effect modifiers with high power. Power is lower for *X*_3_, whose contribution cos^2^(*X*_3_) to *τ* (***X***) has the smallest variance. For the null covariates uncorrelated with the modifiers, the type I error rate stays near the nominal level. For *X*_6_ to *X*_10_, correlated with the modifiers under *ρ* = 0.3, the rate is slightly inflated but remains acceptable. The screen therefore provides calibrated evidence for each covariate, which supports its use in the UNOS analysis.

**Table 4:** Power and type I error at the nominal *α* = 0.05 of the covariate-specific tests under the registry design (*n* = 2000, *p* = 20, *σ* = 3) at block correlation *ρ* = 0 and 0.3 of *X*_1_–*X*_10_. Power is reported for the effect modifiers *X*_1_–*X*_5_ and type I error for the null covariates *X*_6_–*X*_20_.

| **Covariate** | $\rho = 0$ | | $\rho = 0.3$ | |
| --- | --- | --- | --- | --- |
|  | Power | Type I error | Power | Type I error |
| $X_1$ | 0.867 | — | 0.911 | — |
| $X_2$ | 0.841 | — | 0.898 | — |
| $X_3$ | 0.463 | — | 0.515 | — |
| $X_4$ | 0.964 | — | 0.845 | — |
| $X_5$ | 0.925 | — | 0.841 | — |
| $X_6$ | — | 0.038 | — | 0.062 |
| $X_7$ | — | 0.045 | — | 0.057 |
| $X_8$ | — | 0.041 | — | 0.075 |
| $X_9$ | — | 0.053 | — | 0.074 |
| $X_{10}$ | — | 0.048 | — | 0.074 |
| $X_{11}$ | — | 0.045 | — | 0.041 |
| $X_{12}$ | — | 0.042 | — | 0.047 |
| $X_{13}$ | — | 0.044 | — | 0.043 |
| $X_{14}$ | — | 0.041 | — | 0.044 |
| $X_{15}$ | — | 0.039 | — | 0.039 |
| $X_{16}$ | — | 0.041 | — | 0.044 |
| $X_{17}$ | — | 0.047 | — | 0.046 |
| $X_{18}$ | — | 0.043 | — | 0.056 |
| $X_{19}$ | — | 0.049 | — | 0.048 |
| $X_{20}$ | — | 0.040 | — | 0.045 |

### 5.2 Performance on estimating HTE

Beyond detecting heterogeneity, an applied analysis must quantify the benefit for different covariate profiles. We therefore benchmark the *deepHTL* estimator of Section 3.2 against a range of competing estimators (Wager and Athey, 2018; Athey et al., 2019; Künzel et al., 2019; Kennedy, 2023; Chipman et al., 2010). *deepHTL* is among the most accurate estimators across the registry-like settings (Tables S3 and S4). The revision acts as a safeguard in estimation.

## 6 Discussion

This paper illustrates a gatekept strategy for HTE analysis in observational registries, motivated by the choice between BLT and SLT. A positive average treatment effect does not by itself justify individualized decisions, and neither do flexible estimates of individualized effects, unless their variation is first shown to exceed what sampling variability and nuisance estimation error alone would produce. *deepHTL* addresses this problem by combining a global test for heterogeneity, a covariate-specific screen and estimation of individualized and group average effects in one workflow, in which the screen and the estimates are interpreted only after the global test supports heterogeneity.

In the UNOS cohort, the estimated benefit of BLT in post-transplant FEV1% was positive on average but far from uniform across recipients. The largest gains occurred among recipients with low baseline FEV1%, low lung allocation score and low BMI, whereas recipients with preserved baseline lung function gained little. These findings refine the interpretation of the average effect of BLT and suggest that its additional functional benefit is concentrated in clinically identifiable recipient profiles.

The simulations highlight two findings relevant to comparative effectiveness studies of registry data. First, a large constant treatment effect relative to noise can impair outcome fitting, and the resulting error can inflate both global tests. Removing the constant before fitting the nuisance functions improves calibration. Second, both global tests detect heterogeneity with high power, and the covariate-specific screen recovers the true effect modifiers while keeping the type I error for null covariates near the nominal level. The screen thus offers a calibrated alternative to inspecting individualized effect estimates or choosing subgroups after the results are seen.

For applied analysts we recommend the following sequence. Define the causal estimand, the adjustment set and the candidate modifiers before the analysis. Adjust for confounding with flexible cross-fitted nuisance models. Test the global null hypothesis of no heterogeneity before interpreting any individualized effect. If that test rejects, screen the candidate covariates to identify credible effect modifiers. Finally, report group average effects at clinically meaningful thresholds, and keep the statistical evidence for heterogeneity separate from any treatment recommendation. The results of this sequence complement clinical evidence rather than constitute a clinical decision rule, and the sequence applies equally to observational studies outside medicine.

Several limitations remain. The cohort spans 2005–2011, and subsequent changes in allocation policy, surgical practice, perioperative management and recipient selection may limit transportability to current practice. The analysis addresses a single outcome, FEV1%, whereas transplant decisions also depend on survival, perioperative morbidity, quality of life, waitlist consequences and donor organ availability. Measured donor and recipient characteristics are adjusted for, but unmeasured or unadjusted center practice, frailty, disease phenotype or treatment selection factors may still confound the estimated effects. Because the conditional exchangeability assumption cannot be verified from registry data, a sensitivity analysis for conditional treatment effects under unmeasured confounding (Yadlowsky et al., 2022) would complement the estimates reported here.

Finally, the covariate-specific screen showed at most slight type I error inflation at the correlation levels considered, but stronger correlation, as between recipient height and sex in the UNOS cohort, may make it harder to attribute heterogeneity to a single covariate (Hooker et al., 2021).

In summary, *deepHTL* offers a practical test-before-estimate framework for observational HTE analysis. In the UNOS registry it rejects a constant effect of BLT on post-transplant FEV1% and identifies the recipient profiles with the largest estimated benefit. More broadly, the study shows how formal testing for heterogeneity, screening of effect modifiers and clinically interpretable effect summaries can strengthen the analysis of individualized treatment effects in observational registries.

## Acknowledgments

The data reported here have been supplied by the United Network for Organ Sharing as the contractor for the Organ Procurement and Transplantation Network. The interpretation and reporting of these data are the responsibility of the author(s) and in no way should be seen as an official policy of or interpretation by the OPTN or the U.S. Government. This work was partially supported by the National Institutes of Health (R01LM014407 and 1R01HL173044).

## Conflict of interest

The authors declare no potential conflicts of interest.

## Data availability statement

The registry data were provided by UNOS under a data use agreement and cannot be redistributed, and access requests should be directed to UNOS. Code implementing the methods and the simulation studies is available at https://github.com/BZou-lab/deepHTL.

## Supplementary Material

This Supplementary Material contains Appendices A to C, Tables S1 to S4 and Figure S1.

## Appendix A: Algorithms

### Algorithm A1.

Estimation

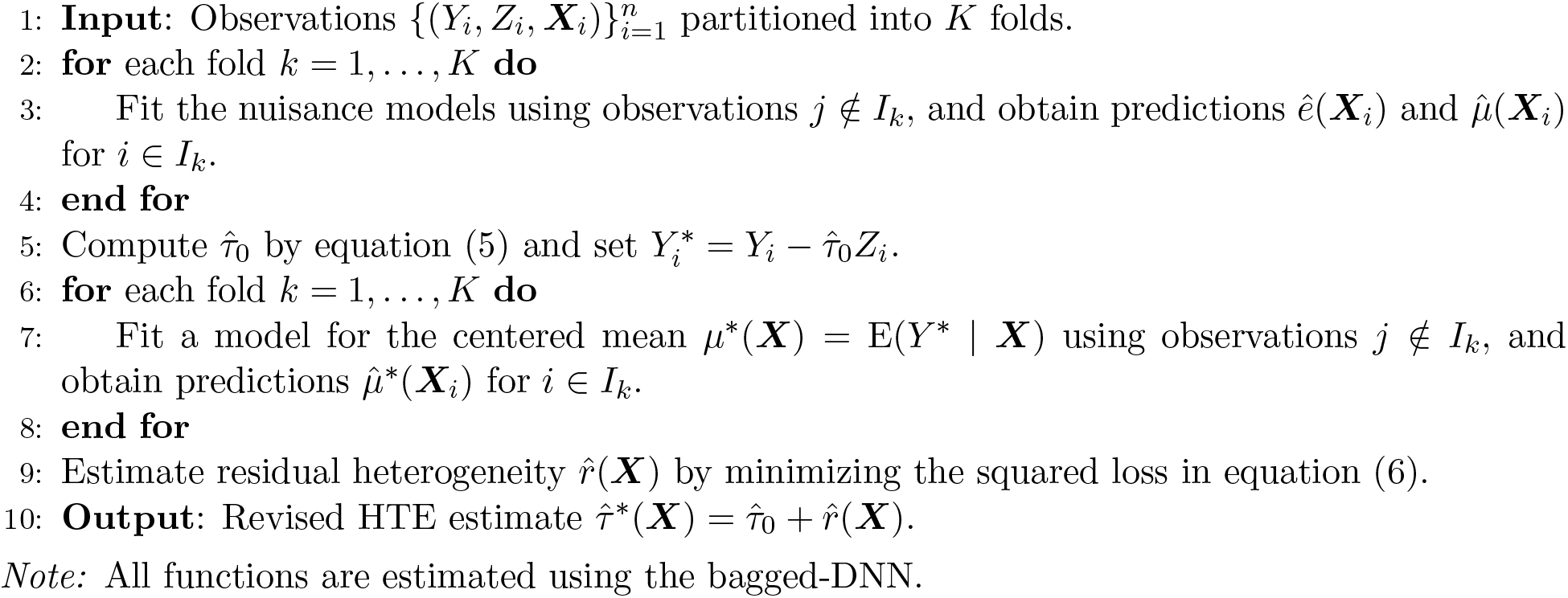

### Algorithm A2.

Testing

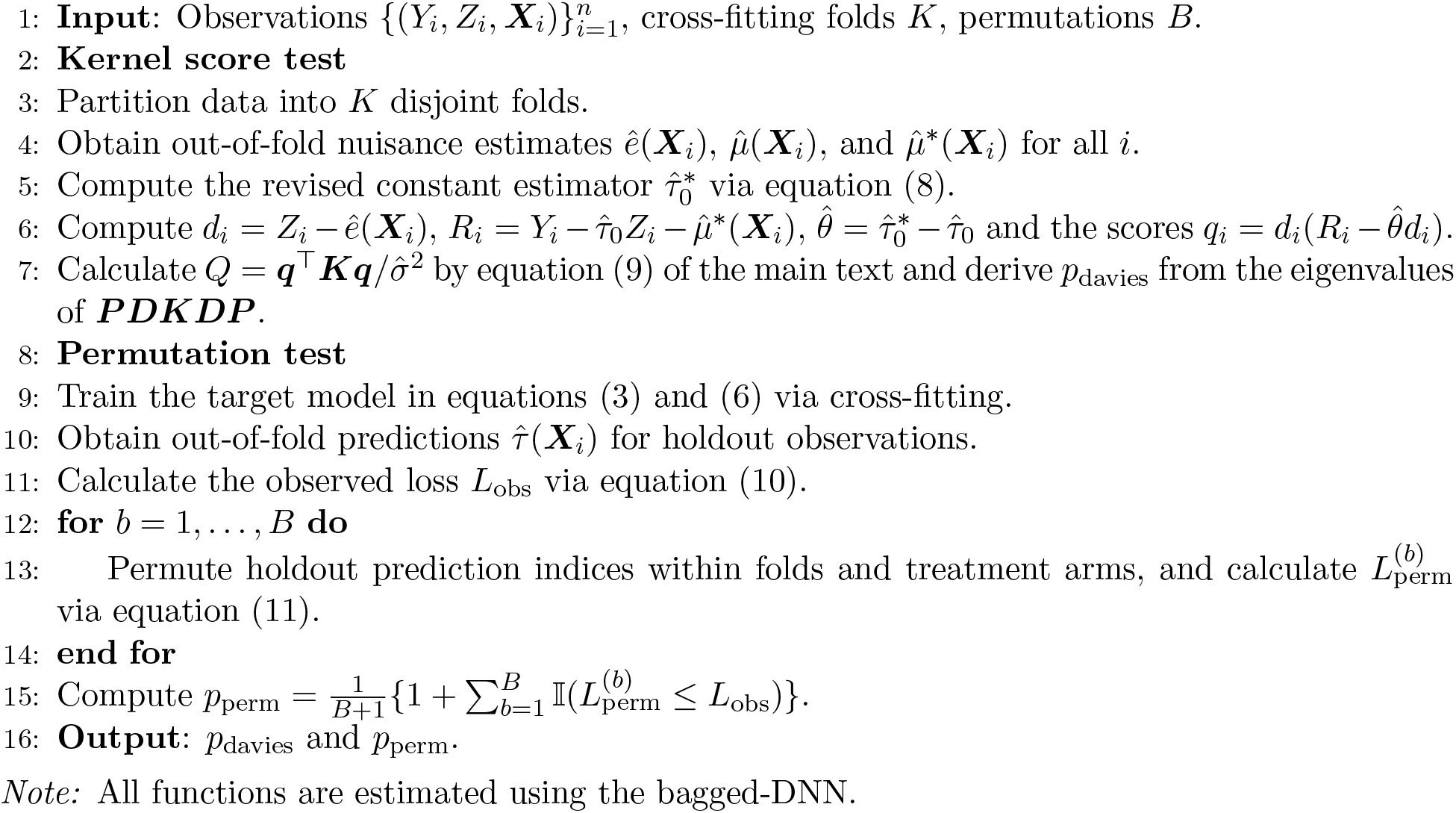

## Appendix B: Implementation settings

### Bagged-DNN (deepHTL and R-DNN)

Nuisance models are cross-fitted over five folds stratified by treatment. The outcome networks also take as inputs the squares and pairwise products of the covariates selected by a lasso with ten-fold cross-validation. Each bagged-DNN has 30 networks with 128, 64 and 32 ReLU units, trained by Adam (learning rate 10^−3^) on bootstrap samples with standardized inputs and outputs, keeping the best out-of-bag epoch. The *L*_1_ penalty is 10^−3^, the outcome models use mini-batches of 128, at most 500 epochs and patience 50, the propensity model 256, 120 and 20, and *ê* is clipped to [0.05, 0.95]. In the UNOS analysis the average treatment effect and GATE are the overall and subgroup means of the augmented inverse probability weighted pseudo-outcomes (Bang and Robins, 2005).

### Other estimators

Lasso: glmnet, penalty by ten-fold cross-validation. XGBoost: the best of ten random draws of the tuning parameters, including the learning rate (0.005 to 0.2) and tree depth (3 to 20), by five-fold cross-validation, up to 1000 rounds and early stopping after ten. KRR: Gaussian kernel, bandwidth (0.5, 1 or 2 times the median pairwise distance) and ridge penalty (13 values from 10^−4^ to 10^2^) by five-fold cross-validation. Causal forest: 2000 trees, all parameters tuned automatically. T-, S-, X- and DR-learners: random forests with 500 trees and minimum node size five, five-fold cross-fitting for the DR-learner. BART T-learner: default prior, one fit per arm. All analyses use *B* = 2000 permutations and 1000 replicates per configuration unless stated otherwise.

## Appendix C: Residual scores of the kernel score test

Let *d*_*i*_ = *Z*_*i*_ − *ê* (***X***_*i*_) and let 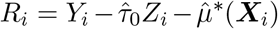 for the revised test and 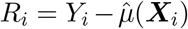 for the unrevised test. Write ***R*** = (*R*_1_, … , *R*_*n*_) , ***d*** = (*d*_1_, … , *d*_*n*_) , ***D*** = diag(***d***) and ***q*** = (*q*_1_, … , *q*_*n*_). Define

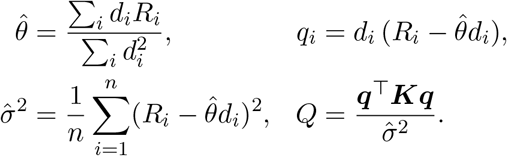

Here 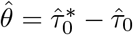 for the revised residual and 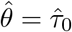 for the unrevised residual.

### Null distribution

Let ***P*** = ***I***_*n*_ − ***dd***^⊤^*/*(***d***^⊤^***d***) be the projection matrix onto the space orthogonal to ***d***. Then 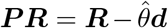 removes the fitted constant effect component, and ***q*** = ***DP R***. Under *H*_0_, a conditionally homoskedastic Gaussian working model motivates the quadratic-form approximation (Liu et al., 2007; Lin et al., 2013)

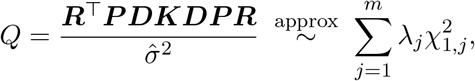

where *λ*_1_, … , *λ*_*m*_ are the nonzero eigenvalues of ***P DKDP*** and the 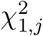 are independent. Davies’ (1980) method computes the upper-tail probability of this mixture.

### Estimated nuisance functions

For the unrevised residual, write *δ*_*e*_ = *ê* − *e* and 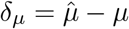. Under *H*_0_, *R*_*i*_ − *τ*_0_*d*_*i*_ = *ε*_*i*_ − *δ*_*µ*_(***X***_*i*_) + *τ*_0_*δ*_*e*_(***X***_*i*_), so

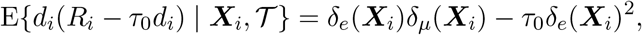

where *T* denotes the training data outside observation *i*’s fold. The conditional mean bias is thus second-order in nuisance errors, involving only products and squares (Chernozhukov et al., 2018). By Cauchy–Schwarz, it is *o*_*p*_(*n*^−1*/*2^) in *L*_1_(*P*_*X*_) if, in each fold, 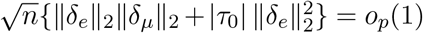, where ∥ · ∥_2_ is the *L*_2_(*P*_*X*_) norm. For fixed *c*, the revised version replaces *δ*_*µ*_ By 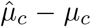, where *µ*_*c*_(***X***) = E(*Y* − *cZ* | ***X***), and *τ*_0_ by *τ*_0_ − *c*. Accurate centering can reduce the squared propensity-error term, and fitting 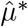 without the large constant can reduce the outcome-model error when |*τ*_0_| is large.

**Table S1:** The 26 adjustment covariates retained by the GL-DR selection procedure, by transplant type.

|  | **BLT** | **SLT** |
| --- | --- | --- |
| | ($n = 9678$) | ($n = 4518$) |
| Recipient age (years), mean (SD) | 53.4 (13.6) | 63.4 (7.5) |
| Recipient BMI, mean (SD) | 24.8 (4.7) | 26.3 (4.1) |
| Lung allocation score, mean (SD) | 40.8 (13.5) | 40.3 (11.1) |
| Recipient diabetes, $n$ (%) | 1985 (20.5%) | 831 (18.4%) |
| Baseline FEV1%, mean (SD) | 37.1 (20.7) | 43.6 (20.4) |
| Recipient sex, female, $n$ (%) | 4049 (41.8%) | 1601 (35.4%) |
| Recipient height (cm), mean (SD) | 170.0 (9.9) | 170.5 (9.6) |
| Recipient Hispanic ethnicity, $n$ (%) | 679 (7.0%) | 303 (6.7%) |
| Oxygen requirement at rest (L/min), mean (SD) | 5.6 (5.3) | 4.7 (4.3) |
| Ventilator support, $n$ (%) | 461 (4.8%) | 71 (1.6%) |
| Six-minute walk distance (ft), mean (SD) | 773.8 (425.8) | 795.0 (411.3) |
| Recipient PO<sub>2</sub> (mmHg), mean (SD) | 76.1 (40.1) | 75.6 (34.0) |
| Donor age (years), mean (SD) | 34.6 (14.0) | 34.9 (13.8) |
| Donor sex, female, $n$ (%) | 3830 (39.6%) | 1671 (37.0%) |
| Donor height (cm), mean (SD) | 171.7 (10.2) | 172.3 (9.9) |
| Donor BMI, mean (SD) | 26.2 (5.5) | 26.2 (5.3) |
| Donor race, Black, $n$ (%) | 1784 (18.4%) | 853 (18.9%) |
| Donor diabetes, $n$ (%) | 698 (7.2%) | 314 (6.9%) |
| Donor CMV serology, positive, $n$ (%) | 6061 (62.6%) | 2782 (61.6%) |
| Donor cause of death, traumatic brain injury (TBI), $n$ (%) | 4199 (43.4%) | 1993 (44.1%) |
| Local or regional allocation, $n$ (%) | 6862 (70.9%) | 2926 (64.8%) |
| Donor to recipient distance (nautical miles), mean (SD) | 204.3 (246.7) | 216.8 (252.4) |
| Ischemic time (h), mean (SD) | 5.6 (1.8) | 4.3 (1.5) |
| Sex match, $n$ (%) | 6649 (68.7%) | 3126 (69.2%) |
| Race match, $n$ (%) | 5448 (56.3%) | 2533 (56.1%) |
| Time to follow-up FEV1 (days), mean (SD) | 377.4 (62.9) | 380.9 (70.9) |

**Table S2:**
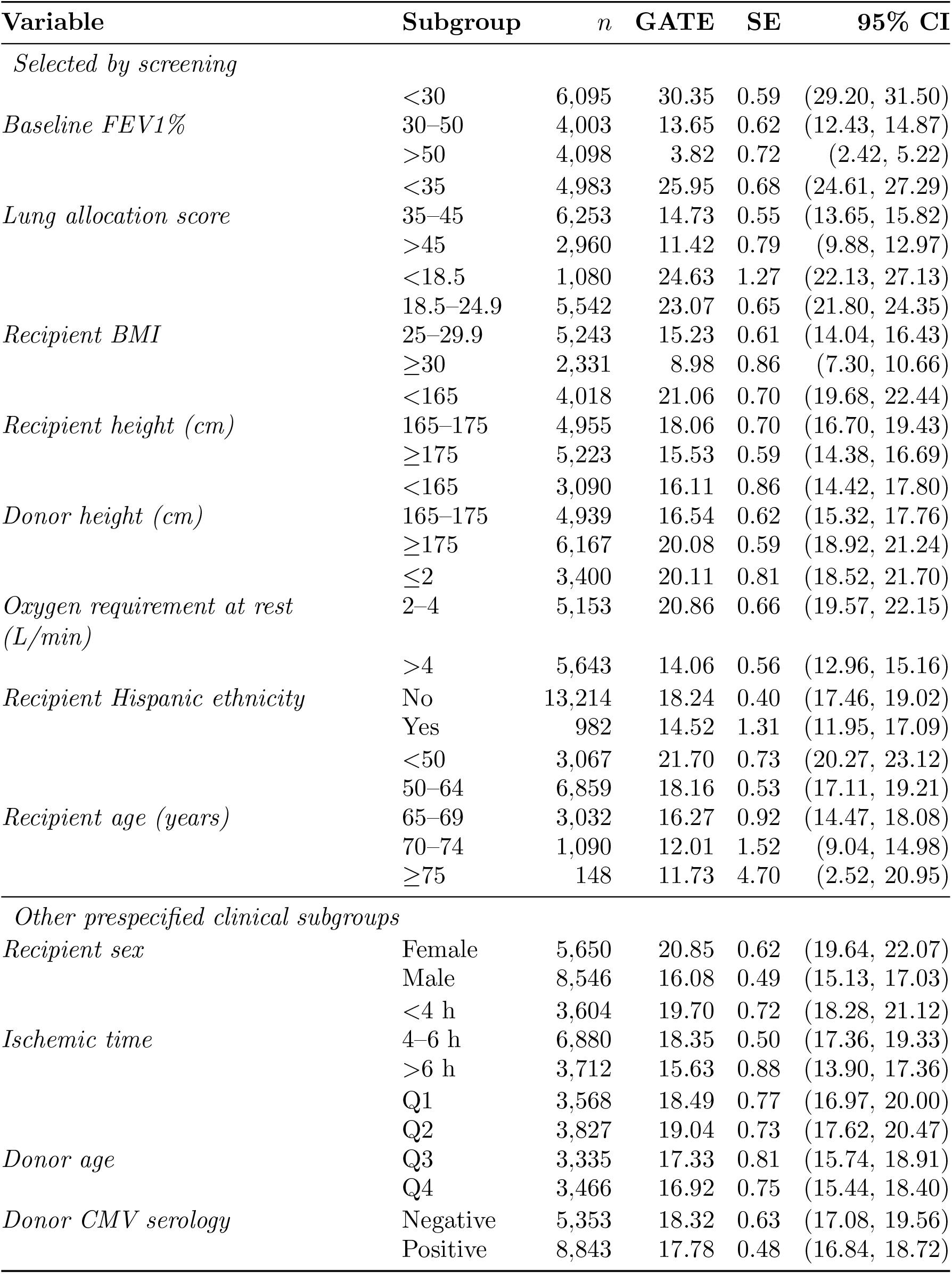
Group average treatment effects (GATE, BLT versus SLT) on post-transplant FEV1% for the screen-selected variables and prespecified clinical subgroups.

**Table S3:** Mean log-MSE of all estimators over 1000 replicates for the HTE effect function at *ρ* = 0. Each estimator is trained on one data set and evaluated on an independent test set of the same size, and log-MSE is the logarithm of the mean squared error of 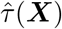 over the test set. Monte Carlo standard errors of all entries are below 0.01. *R-* and *Rev-* denote the unrevised and revised R-learner with the given base learner, Lasso, XGBoost or kernel ridge regression (KRR), and R-DNN is the unrevised counterpart of *deepHTL*. The established estimators are the causal forest, the X-, T- and S-learners, the DR-learner and a T-learner with Bayesian additive regression tree base learners (BART-T).

| $\tau(\mathbf{X}) = -1 + X_1X_2 + \cos^2(X_3) + \max(X_4 - X_5, 0)$ | | | | |
| --- | --- | --- | --- | --- |
| $n = 1000$ | | | | |
| Method | $p = 20, \sigma = 1$ | $p = 40, \sigma = 1$ | $p = 20, \sigma = 3$ | $p = 40, \sigma = 3$ |
| deepHTL | -0.484 | -0.108 | 0.418 | 0.558 |
| R-DNN | -0.483 | -0.103 | 0.417 | 0.548 |
| R-Lasso | 0.305 | 0.332 | 0.484 | 0.528 |
| Rev-Lasso | 0.307 | 0.335 | 0.480 | 0.525 |
| R-XGBoost | 0.146 | 0.293 | 0.548 | 0.596 |
| Rev-XGBoost | 0.152 | 0.310 | 0.554 | 0.602 |
| R-KRR | -0.183 | 0.376 | 0.496 | 0.574 |
| Rev-KRR | -0.165 | 0.375 | 0.506 | 0.581 |
| BART-T | -0.275 | -0.017 | 0.887 | 1.047 |
| DR-learner | 0.327 | 0.412 | 0.808 | 0.772 |
| T-learner | 0.213 | 0.328 | 0.606 | 0.622 |
| X-learner | 0.233 | 0.349 | 0.446 | 0.509 |
| Causal forest | 0.215 | 0.337 | 0.444 | 0.526 |
| S-learner | 0.437 | 0.537 | 0.550 | 0.606 |
| $n = 2000$ | | | | |
| Method | $p = 20, \sigma = 1$ | $p = 40, \sigma = 1$ | $p = 20, \sigma = 3$ | $p = 40, \sigma = 3$ |
| deepHTL | -0.637 | -0.384 | 0.269 | 0.318 |
| R-DNN | -0.638 | -0.398 | 0.271 | 0.318 |
| R-Lasso | 0.247 | 0.260 | 0.369 | 0.409 |
| Rev-Lasso | 0.245 | 0.260 | 0.365 | 0.410 |
| R-XGBoost | -0.185 | 0.083 | 0.443 | 0.505 |
| Rev-XGBoost | -0.146 | 0.083 | 0.444 | 0.500 |
| R-KRR | -0.668 | 0.146 | 0.324 | 0.498 |
| Rev-KRR | -0.682 | 0.140 | 0.329 | 0.503 |
| BART-T | -0.660 | -0.508 | 0.533 | 0.688 |
| DR-learner | 0.209 | 0.285 | 0.619 | 0.602 |
| T-learner | 0.060 | 0.191 | 0.439 | 0.464 |
| X-learner | 0.136 | 0.257 | 0.330 | 0.405 |
| Causal forest | 0.096 | 0.220 | 0.346 | 0.442 |
| S-learner | 0.328 | 0.446 | 0.499 | 0.573 |

**Table S4:** Mean log-MSE of all estimators over 1000 replicates for the HTE effect function at *ρ* = 0.3. Monte Carlo standard errors of all entries are below 0.01. Notation as in Table S3.

| $\tau(\mathbf{X}) = -1 + X_1X_2 + \cos^2(X_3) + \max(X_4 - X_5, 0)$ | | | | |
| --- | --- | --- | --- | --- |
| $n = 1000$ | | | | |
| Method | $p = 20, \sigma = 1$ | $p = 40, \sigma = 1$ | $p = 20, \sigma = 3$ | $p = 40, \sigma = 3$ |
| deepHTL | -0.443 | -0.132 | 0.358 | 0.499 |
| R-DNN | -0.440 | -0.128 | 0.362 | 0.499 |
| R-Lasso | 0.327 | 0.360 | 0.491 | 0.527 |
| Rev-Lasso | 0.326 | 0.361 | 0.491 | 0.527 |
| R-XGBoost | -0.003 | 0.135 | 0.502 | 0.561 |
| Rev-XGBoost | 0.013 | 0.153 | 0.504 | 0.560 |
| R-KRR | -0.215 | 0.315 | 0.462 | 0.544 |
| Rev-KRR | -0.228 | 0.287 | 0.439 | 0.532 |
| BART-T | -0.280 | -0.058 | 0.816 | 0.962 |
| DR-learner | 0.160 | 0.277 | 0.749 | 0.696 |
| T-learner | 0.022 | 0.151 | 0.499 | 0.513 |
| X-learner | 0.091 | 0.222 | 0.354 | 0.419 |
| Causal forest | 0.147 | 0.291 | 0.405 | 0.480 |
| S-learner | 0.354 | 0.458 | 0.527 | 0.576 |
| $n = 2000$ | | | | |
| Method | $p = 20, \sigma = 1$ | $p = 40, \sigma = 1$ | $p = 20, \sigma = 3$ | $p = 40, \sigma = 3$ |
| deepHTL | -0.615 | -0.359 | 0.124 | 0.228 |
| R-DNN | -0.612 | -0.355 | 0.126 | 0.231 |
| R-Lasso | 0.267 | 0.286 | 0.402 | 0.430 |
| Rev-Lasso | 0.268 | 0.287 | 0.400 | 0.436 |
| R-XGBoost | -0.374 | -0.197 | 0.339 | 0.430 |
| Rev-XGBoost | -0.353 | -0.171 | 0.350 | 0.433 |
| R-KRR | -0.711 | -0.009 | 0.243 | 0.467 |
| Rev-KRR | -0.717 | -0.006 | 0.228 | 0.447 |
| BART-T | -0.737 | -0.525 | 0.474 | 0.658 |
| DR-learner | -0.055 | 0.116 | 0.509 | 0.516 |
| T-learner | -0.199 | -0.024 | 0.273 | 0.329 |
| X-learner | -0.062 | 0.101 | 0.181 | 0.294 |
| Causal forest | -0.089 | 0.101 | 0.283 | 0.411 |
| S-learner | 0.166 | 0.320 | 0.414 | 0.527 |

**Figure S1:**
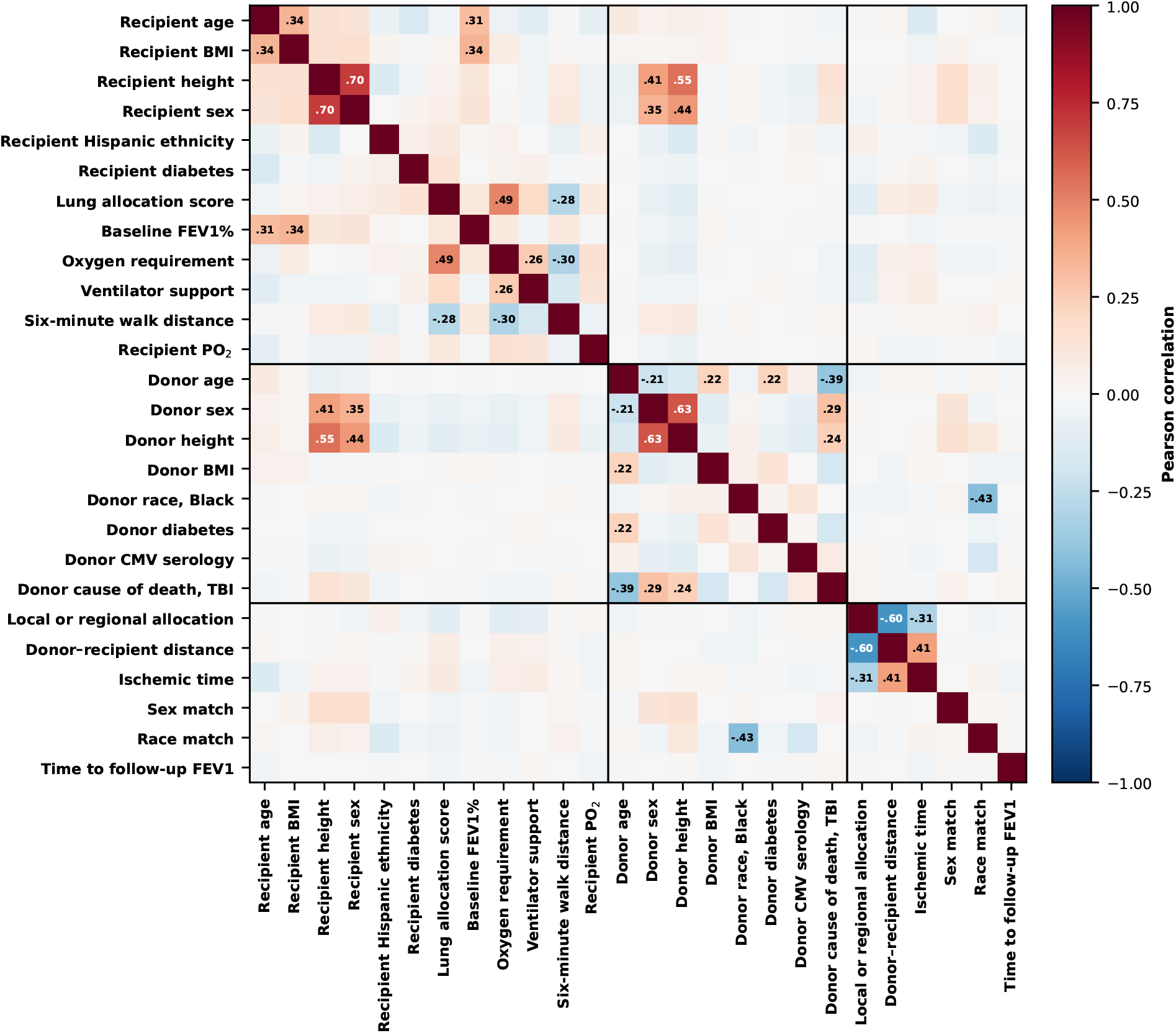
Heat map of pairwise Pearson correlations among 26 covariates in the UNOS cohort.

## Notes

https://github.com/BZou-lab/deepHTL

### Competing Interest Statement

The authors have declared no competing interest.

### Summary of Updates

Add a screening step; add more real data analysis; recontsruct the paper

